# Optical Control of Invertebrate nAChR and Behaviors with Dithienylethene-Imidacloprid

**DOI:** 10.1101/2021.08.27.457891

**Authors:** Chao Zhang, Qi Xu, Zhiping Xu, Long Wang, Zewen Liu, Zhong Li, Xusheng Shao

## Abstract

Photopharmacology has changed established methods of studying receptor functions, allowing for increasing spatiotemporal resolution. However, no photopharmacological tools are available for the invertebrate nicotinic acetylcholine receptor (nAChR). Here, we report a photochromic ligand, dithienylethene-imidacloprid (DitIMI), targeting invertebrate nAChR. We demonstrated that DitIMI has low spontaneous *in vivo* and *in vitro* activity but can be photoisomerized to a highly active closed-form. This photoisomerization can further be translated to photomodulation of neuron membrane potential and behavioral responses of living mosquito larvae and American cockroaches. Furthermore, we discovered that DitIMI is a specific reporter for fluorescence polarization based high-throughput screening of nAChR ligands.

## INTRODUCTION

Nicotinic acetylcholine receptors (nAChRs) are pentameric ligand-gated transmembrane ion channels that mediate synaptic transmission and information processing with binding sites for endogenous acetylcholine and exogenous ligands (Taly et al., 2009; Bouzat and Sine, 2018). Hypofunction of nAChRs is associated with many neurological diseases that influence tens of millions of people, such as Alzheimer′s and Parkinson′s disease, epilepsy, and schizophrenia (Hurst et al., 2013). nAChRs are also the targets of neonicotinoid, nereistoxin and spinosad pesticides whose annual sale volume approaches three billion US dollars (Casida, 2018). The critical role of nAChRs stimulated isolation and preparation of a diverse array of natural or synthetic ligands, e.g., subtype-selective (Mazurov et al., 2011), radioisotope labelled (Liu et al., 2013), photoaffinity (Hamouda et al., 2014) and fluorescent ligands (Grandl et al., 2007; Jing et al., 2018). These ligands are basic instruments for identification, pharmacological characterization and functional modulation of nAChRs. However, the control offered by conventional ligands lacks high spatiotemporal resolution. This shortcoming makes it difficult to elucidate the timing and location of ligand-receptor interactions.

The emergence of photopharmacology provides a solution for the limitations of conventional ligands (Hüll et al., 2018; Lerch et al., 2016; Broichhagen et al., 2015; Velema et al., 2014). This innovative technology uses photoresponsive ligands to realize spatiotemporal control over a variety of biological processes under the aid of light (Hoorens and Szymanski, 2018; Berlin and Isacoff, 2017; Kienzler and Isacoff, 2017; Bregestovski et al., 2018; van Leeuwen et al., 2017; Gautier et al., 2014). The most commonly-used photopharmacological tools are photochromic ligands (PCLs) and photochromic tethered ligands (PTLs) (Hüll et al., 2018; Kramer et al., 2013). PCLs and PTLs contain a photoswitch whose structure can be changed reversibly upon irradiation, along with the alternation of binding circumstances. The advantage of not requiring genetic modification makes PCL a popular and convenient strategy with broad applications in many biologically-important targets, such as ionotropic or metabotropic glutamate receptors (iGlu or mGlu) (Reiner et al., 2015; Goudet et al., 2018; Pittolo et al., 2014), NMDA receptors (Berlin et al., 2016), GABA receptors (Yue et al., 2012; Lutz et al., 2018; Stein et al., 2012), protein kinase C (Frank et al., 2016), epithelial sodium channel (Schönberger et al., 2014), L-type Ca^2+^ channels (Fehrentz et al., 2018), vanilloid receptor (Frank et al., 2015), microtubule (Borowiak et al., 2015), transient receptor (TRP) channels (Leinders-Zufall et al., 2018; Lichtenegger et al., 2018), glycine receptors (Gomila et al., 2020), cannabinoid receptor (Westphal et al., 2017) and potassium channel (Barber et al., 2016; Banghart et al., 2009) as well as nAChRs (Tochitsky et al., 2012; Damijonaitis et al., 2015; Damijonaitis et al.; Deal et al., 1969; Kagabu et al., 2010).

Photopharmacology studies of nAChRs advance our understanding of their roles in the nervous system. An array of nAChR PCLs or PTLs have been developed based on known agonists or antagonists since photochromic antagonist *p*-phenylazophenyltrimethylammonium was first reported in 1969 (Deal et al., 1969). However, recent progress has been slow: only one PCL for neuronal α7 nAChR (AzoCholine) and two PTLs (maleimide-azobenzene-Ach and maleimide-azobenzene-homocholine, respectively) for α3β4 and α4β2 nAChR have been developed. nAChRs are pentameric receptors composed of various permutations of subunits, and this subtype diversity requires development of more selective PCLs. However, to date, all photopharmacological ligands have been designed for vertebrate nAChRs, and lack the counterpart acting on invertebrate nAChR. All nAChR PCLs and PTLs are derived from an azobenzene photoswitch, while the utility of other types of photoswitches is unknown. Furthermore, details of the potential use of nAChR PCLs to control the behavioral response of living organisms remain elusive. We present a dithienylethene (Dit)-based nAChR PCL, named dithienylethene-imidacloprid (DitIMI). Through a combination of chemical and biological techniques, we demonstrate that DitIMI enables optical control of invertebrate nAChR, DUM neurons and behavioral responses of mosquito larvae and American cockroaches. DitIMI also enables high-throughput nAChR ligand screening based on fluorescence polarization (FP).

## RESULTS

### DitIMI is generated by linking two imidacloprid molecules with dithienylethene

PCLs are usually derived from subtle linkage of a photoswitch to the core of a high-potency ligand. Imidacloprid (IMI) is a selective nAChR agonist whose sales volume approaches one billion USD annually (Casida, 2018). Kagabu *et al.* prepared a set of divalent IMI analogues with varied length linkers and verified heptamethylene-tethered IMI (HepIMI) as the most potent ligand (Kagabu et al., 2010). The linker length is crucial to the activity and the optimal distance between two nitrogen atoms is around 9.96 Å (Fig. 1A, HepIMI). By replacing heptamethylene with azobenzene, we previously prepared a photoswitchable AzoIMI whose activity increased 5-fold during *trans*-to-*cis* isomerization accompanying N-N distance change from 12.59 Å to 8.14 Å (Fig. 1A) (Xu et al., 2015). However, the relatively low activity of AzoIMI limits its further applications, probably due to the poor solubility of azobenzene (Beharry and Woolley, 2011).

**Figure 1.**
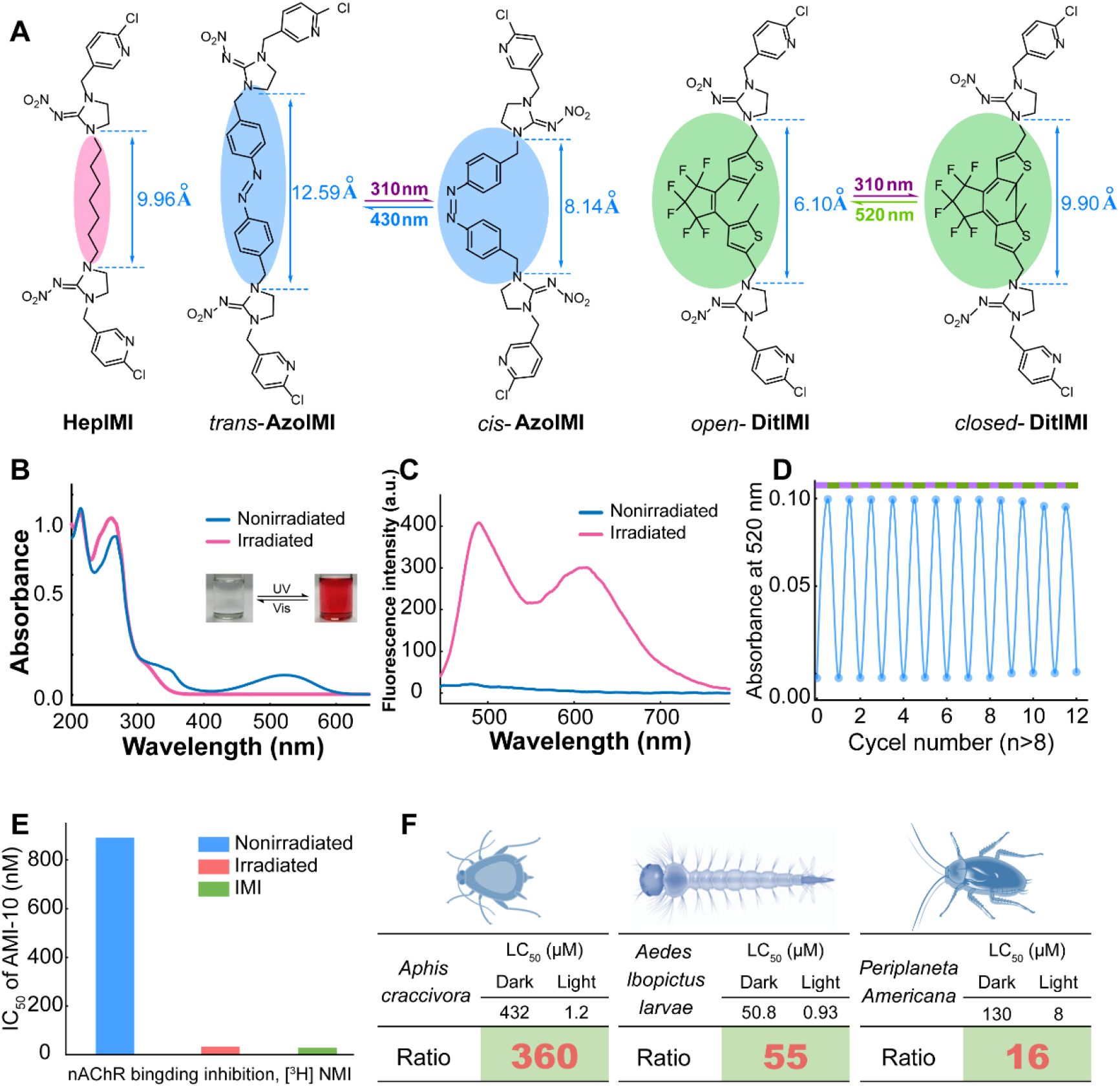
Molecular design, photophysiochemical property characterization and *in vitro* and *in vivo* activity. (A) Molecular design of nAChR PCL DitIMI. (B) UV-Vis absorption spectra of DitIMI (44 μM in acetonitrile) before and after irradiation at 320 nm, inset pictures are color changes of DitIMI solution (1 mM in acetonitrile). (C) Fluorescence emission spectra of open- and closed-DitIMI (44 μM in acetonitrile). (D) Reversible photostability evaluation of DitIMI upon irradiation alternately with UV and visible light (520 nm). (E) Binding affinity of DitIMI and IMI to housefly (*Musca domestica*) using a tritium labelled nitromethylene analogue of IMI ([^3^H]NMI) as a radiotracer. (F) *In vivo* activity of DitIMI against cowpea aphids (*Aphis craccivora*), mosquito larvae (*Aedes albopictus*) and cockroaches (*Periplaneta americana*) before and after irradiation.

Based on the above results, we intended to investigate other photoswitchable linkers starting from calculating N-N distance and found that dithienylethene-tethered IMI might be a potential candidate. The N-N distance of open-DitIMI is 6.61 Å, while its photoisomer (closed-DitIMI) has a distance of 9.90 Å, which is close to that of high potency HepIMI. DitIMI was synthesized in seven steps (Supplementary Fig. 1).

### DitIMI exhibits excellent photoisomerization properties

DitIMI exists completely in its thermally stable open-form. Upon irradiation with ultraviolet light (320 nm), open-DitIMI gradually transformed to its closed-form with appearance of an absorption band at 520 nm and red fluorescence (Fig. 1B and 1C). In the photostationary state, 67% closed-DitIMI was produced. The closed-to-open isomerization occurred at the presence of λ = 520 nm light and this process can be triggered back and forth over ten cycles without any obvious fatigue (Fig. 1D). Both photoisomers were stable in water and phosphate-buffered saline (PBS), guaranteeing stability during biological tests.

### Photoisomers show differential *in vitro* and *in vivo* activity

To evaluate the efficacy level and differential activity, housefly (*Musca domestica*) nAChR affinity was evaluated using a tritium labelled nitromethylene analogue of IMI ([^3^H]NMI) as a radiotracer. [^3^H]NMI is a good reporter for affinity to invertebrate nAChRs (Shao et al., 2013). Open-DitIMI showed low *in vitro* binding potency to *Musca* nAChR (IC_50_ = 892 nM). Illumination with UV light significantly enhanced the affinity (IC_50_ = 33.2 nM) close to that of IMI (IC_50_ = 28.8 nM), indicating a 27-fold activity difference (Fig. 1E).

Having identified closed-DitIMI as a high potency ligand in the *Musca* nAChR assay, we then evaluated *in vivo* activity of DitIMI towards three different insect species: cowpea aphids (*Aphis craccivora*), mosquito larvae (*Aedes albopictus*) and American cockroaches (*Periplaneta americana*) (Fig. 1F). Cowpea aphids were used for evaluating potency as IMI is extremely effective against this sucking insect. Transparent mosquito larvae were used for a behavioral study and cockroaches were used for neuron membrane potential studies. Huge, light-induced activity enhancement, approximately 355-fold, was observed with cowpea aphids, indicating almost complete switching on and off of the activity. Similar light-dependent activation was also observed in mosquito larvae and cockroaches, with 22- and 16-fold activity increases, respectively. The photoactivation is probably partly due to the N-N distance change during isomerization. The above *in vitro* and *in vivo* experiments established that DitIMI was almost inactive in its open-form and can be quickly photoisomerized to a highly potent closed-form.

### Reversible photoisomerization occurs in neurons and living mosquito larvae

Following successful identification of activity differences, we next investigated *in vivo* and *in vitro* real-time isomerization ability trigged by light. The self-contained fluorescence of closed-DitIMI makes it a good indicator for tracing isomerization and imaging studies. The freshly-separated dorsal unpaired median (DUM) neuron of *P. americana* was treated with DitIMI (30 μM) and subjected to UV irradiation. Red fluorescence gradually appeared and approached saturation after 120 s of irradiation (Fig. 2A). The disappearance of red fluorescence was achieved when exposed to λ = 520 nm light for 90 s and this process can be modulated for multiple cycles, indicating occurrence of reversible isomerization. A similar trend was also observed in mosquito larvae treated with DitIMI (30 μM). Irradiation of the treated larvae with UV light led to the occurrence of isomerization *in vivo* accompanying the generation of red fluorescence mainly concentrated in the gastric caeca and gut lumen (Fig. 2B). The above two experiments verify that reversible photoisomerization can easily and efficiently occur inside neurons and living organisms.

**Figure 2.**
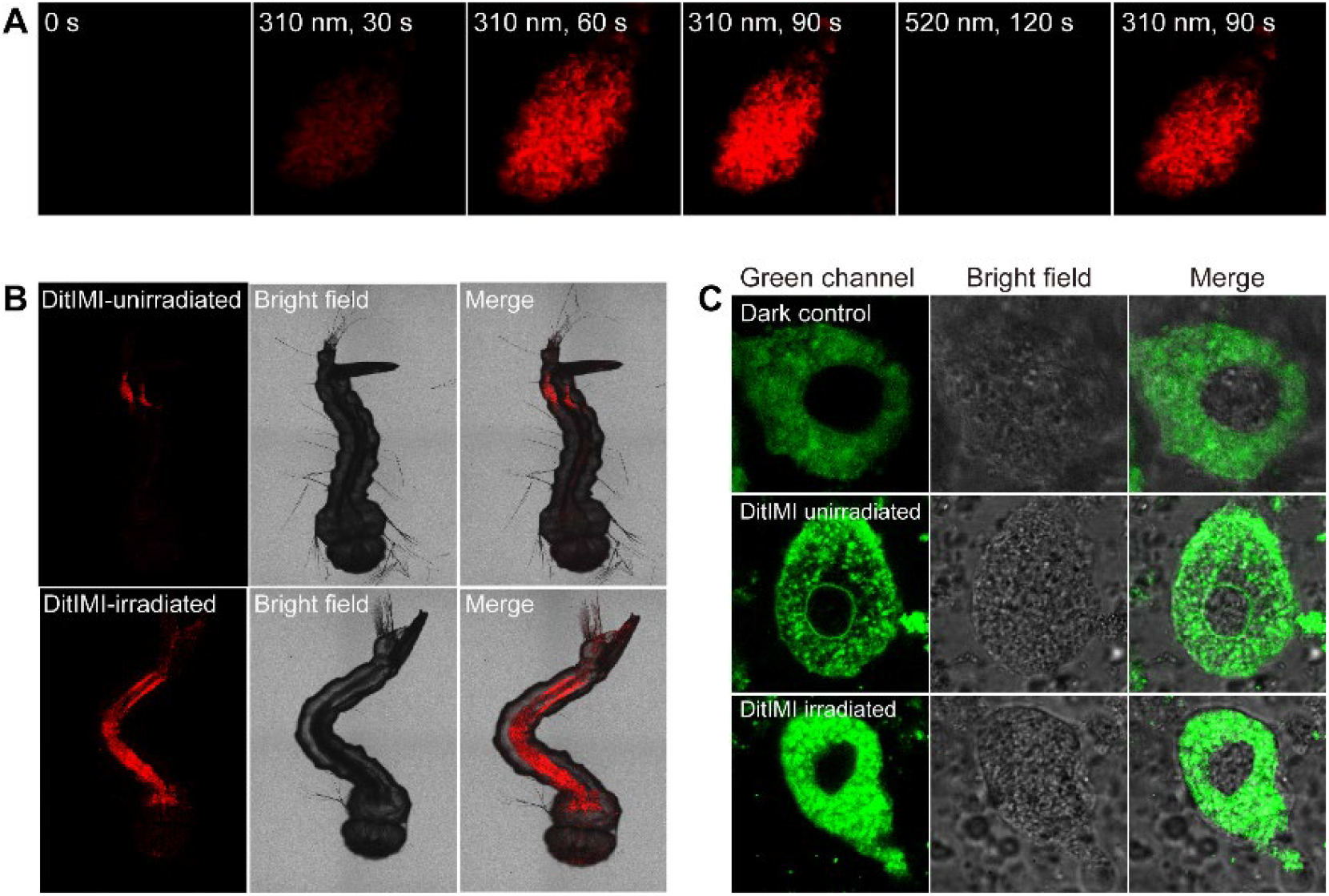
Photoisomerization in nerve cell, living mosquito larvae and cell potential evaluation. (A) Photoconversion of DitIMI (30 μM) in cockroach DUM neurons upon alternate irradiation with UV (320 nm) and visible light (520 nm). (B) Confocal images of living mosquito larvae treated with DitIMI (30 μM) in the dark and upon light irradiation. (C) Membrane potential analysis using DiBAC_4_(3) as a fluorescence staining dye. CK: DUM neuron; unirradiated: neuron treated with DitIMI (30 μM) in the dark; irradiated: neuron treated with DitIMI (30 μM) plus light. Mean fluorescence intensity from top to bottom: 57, 93 and 173.

### DitIMI mediates membrane potential of DUM neurons through perturbing nAChR functions

We next investigated the ability of DitIMI in modulating neurons. We used cockroach DUM neurons because they contain IMI-sensitive nAChR subtypes with DiBAC_4_(3) as the fluorescence dye(Tan et al., 2007). Membrane potential indicator DiBAC_4_(3) is able to enter cells after alternation of membrane polarity with enhanced fluorescence. Light alone does not influence the cell (mean fluorescence intensity 6.93). IMI treated neurons showed fluorescence enhancement in a dosage-dependent manner. Treatment with open-DitIMI had a slight effect on the neurons due to its low affinity to the receptor (mean fluorescence intensity 9.61). Photo illumination induced accumulation of dye inside cells presenting a fluorescence increase of around 2.8-fold (Fig. 2C). This result indicated that IMI induced depolarization of DUM neurons and alternation of membrane potential resulting in an influx of sodium by opening and closing the nAChR channel.

### DitIMI enables optical control of behavior of living mosquito larvae and cockroaches

To examine the possibility of using DitIMI to actively control or predict the behavior of living organisms, we used mosquito larvae and cockroaches to evaluate light-induced behavioral responses (Fig. 3A, 3B). The above observations established that DitIMI acts on nAChRs as an agonist in a light-dependent manner. Interaction with a nAChR agonist can cause behavioral excitement, leading to muscle spasms, jumping and tremors. The transparent feature of mosquito larvae makes light delivery studies simple to conduct. We used a portable UV-lamp as the light source for the initiation of action potential. Light or DitIMI did not solely cause any toxicity to the larvae (Supplementary Video 1). Photostimulation of DitIMI-treated larvae elicited the characteristic increased motility. Cessation of UV light illumination and treatment with 520 nm light led to the recovery of mosquito larvae and this process was reversible for several cycles (Supplementary Video 2). We recorded the moving trial and distance (Fig. 3C), establishing the correlation of nAChR activation with the excited behavioral output.

**Figure 3.**
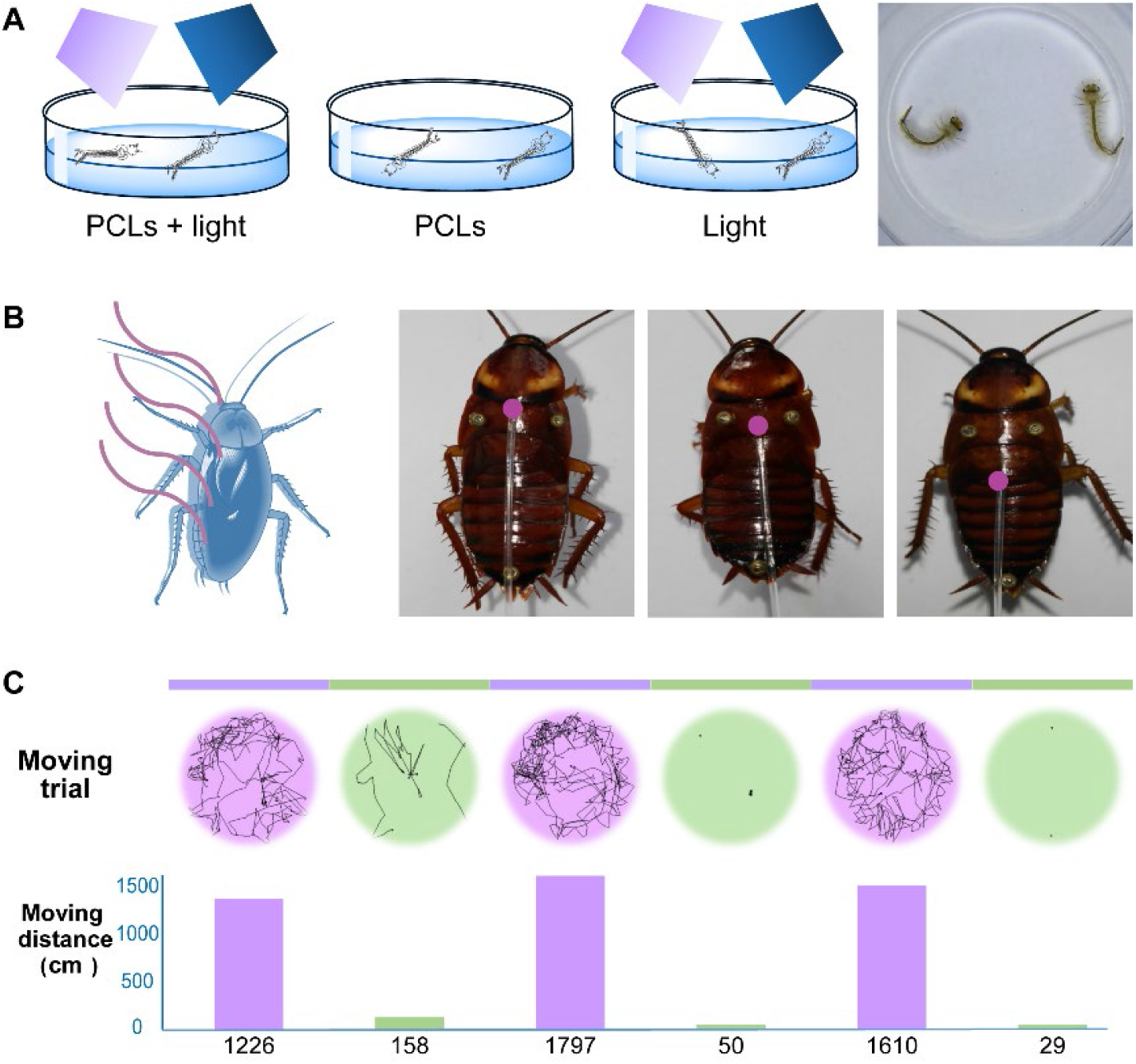
Optical modulation of behaviors of mosquito larvae and cockroaches. (A) Schematic diagram of photomodulation of behaviors of mosquito larvae. (B) Schematic diagram of photomodulation of behaviors of cockroaches treated with DitIMI (12.5 μg/pest) equipped with optical fiber around brain (left), prothoracic ganglion (middle) and mesothoracic ganglion (right). (C) The moving trajectory and distance of mosquito larvae treated with DitIMI (30 μM) at a time interval of 90 s irradiation. The horizontal bar at the top indicates alternative irradiation with UV and visible light (520 nm); the circles in the middle are the movement trajectory of the larvae calculated by ImageJ; the vertical bars at the bottom show the moving distance of mosquito larvae.

Cockroaches are opaque and an optical fiber was therefore used to introduce light inside the body. The brain and vertical nerve cord are the central nervous system (CNS) of the cockroach. We fixed cockroaches with pins to prevent escape. Then an optical fiber was inserted into the position where the CNS was located (Fig. 3B). We first investigated the light-induced behavioral response with the optical fiber placed in the brain. Poisoning signs were monitored thirty minutes after intersegmental membrane injection of DitIMI. Cockroaches loaded with the ligand and light were twittering violently. In comparison, open-DitIMI or light alone did not induce any obvious toxicity to cockroaches (Supplementary Video 3). Illumination of cockroaches harboring DitIMI at the prothoracic or mesothoracic ganglion caused similar excitement effects (Supplementary Videos 4 and 5). Switching to green light did not stop exciting the cockroach, showing irreversible regulation in an opaque organism.

### DitIMI is a good indicator for FP-based high-throughput screening of nAChR ligands

The above results verified that closed-DitIMI not only has high nAChR affinity but also emits fluorescence. These dual features inspired us to examine the possibility of a fluorescence polarization (FP) based assay for nAChR ligands. DitIMI has two maximum emission wavelengths at 490 nm and 610 nm. With establishment of its fluorescence properties, we set out to develop and optimize an FP binding assay using cockroach nAChRs. A saturation binding experiment determined that the dissociation constant (K_d_) of DitIMI is 13.51 nM (Fig. 4A).

**Figure 4.**
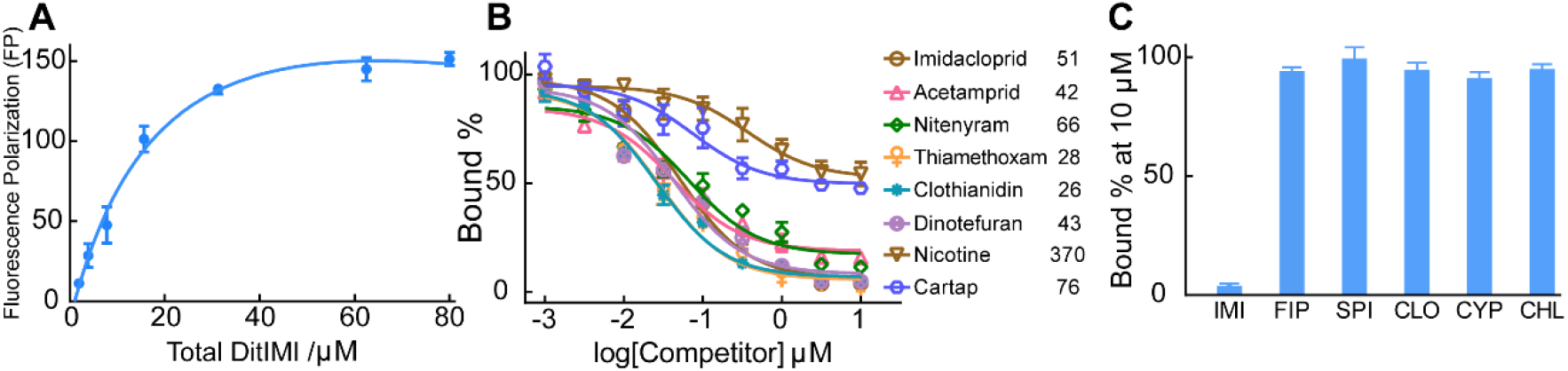
High-throughput nAChR ligands screening using fluorescence polarization. (A) Saturation binding of DitIMI with cockroach nAChR membranes. (B) Competitive binding of various nAChR ligands using DitIMI (40 μM) as a tracer in cockroach membrane protein (227 mg/L). (C) Affinity evaluation of ligands acting on the nervous system.

We then assessed the ability of DitIMI as a tracer for a FP competitive binding assay. The competition experiment was optimized using IMI as a competitive ligand to determine optimal concentrations of protein and DitIMI. A total of 227 mg/L cockroach membrane protein and 40 μM DitIMI provided a good dynamic range for a competition study. A set of known ligands were then screened for inhibiting DitIMI binding with nAChR (Fig. 4B). Neonicotinoids are potent agonists of invertebrate nAChR and can displace DitIMI from protein with IC_50_ values ranging from 20 nm to 70 nm. The less selective ligand nicotine had much lower affinity to cockroach nAChR. The antagonist cartap also displaced DitIMI suggesting that it shares the same binding site with the tracer. To identify the specificity of this method and exclude the possibility that any chemical acting on the nervous system can perturb binding equilibrium, other toxins acting on the nervous system were evaluated (Fig. 4C). Fipronil (GABA antagonist), spinosad (nAChR activator), chlorpyrifos (AChE inhibitor), cypermethrin (sodium channel modulator), and chlorantraniliprole (ryanodine receptor modulator) did not have any influence on DitIMI binding, indicating the robustness of our method for high-throughput screening of ligands acting on nAChR. The activity trend obtained here is consistent with that obtained in a radioligand [3H]HMI binding assay(Shao et al., 2013).

## DISCUSSION

It is now widely appreciated that the ability to use photocontrol in a variety of biological processes can facilitate major advances in understanding of both ligand-receptor and ligand-insect interactions at increasing spatiotemporal resolution. Freely diffusible PCLs acting on native receptors have the benefits of no genetic engineering and easier preparation and use, albeit with less precision. However, progress of nAChR PCL research over the past 30 years has been slow with only one reported example.

We thus established a high-performance dithienylethene-type nAChR PCL, DitIMI, to enable a variety of optical controls from the molecular through to the whole way to the behavioral level. Our PCL has four notable features: it is the first nAChR PCL derived from dithienylethene; the first acting on invertebrate nAChR; the first used for behavioral modulation; and the first ligand for high-throughput screening. DitIMI has high receptor potency and reversible on/off switching of activity, enabling photomodulation of nAChR, which can then be translated to neuronal and behavioral changes. Of note, fast and real-time modulation of the behavioral response of living mosquito larvae and cockroaches was achieved, providing an unprecedented example of a behavioral study using nAChR PCL. Considering the complexity and high cost of using mammalian organisms for research, the outcomes obtained in less complex and comparatively inexpensive invertebrates might provide some clues for the development of new therapeutic technologies.

All previously-described nAChR PCLs are derived from azobenzene photoswitches (Damijonaitis et al., 2015). Azobenzenes are widely used in photopharmacology due to easy attachment, photostability and high transform efficiency (Beharry and Woolley, 2011). However, their rigidity and poor solubility limits biological applications and large activity variations (Zhang and Tian, 2018). We previously investigated azobenzene and azopyridine derived imidacloprid as nAChR PCLs with the best compound showing 5-fold light-induced activity enhancement, but the activity level was relatively low (Xu et al., 2015). Dithienylethene has good thermal stability for both isomers and can only be switched on/off by light. This property guarantees accuracy of biological testing results by excluding interference of thermal isomerization. Our PCL represents the first example of a Dit-based nAChR ligand. The results demonstrated that attachment of Dit is workable in nAChR PCL derivatization, albeit with a relatively small photo-triggered change in molecular geometry.

Many chemical, biological and physical external stimuli are imposed to regulate living organisms leading to different behavioral outcomes, such as body impairment, paralysis, excitement, dysfunction, repellency, growth disorders and eventual death. Intentional control over behaviors through an activity-controllable ligand in a spatiotemporal manner has outstanding merits for understanding insect-stimuli interactions and facilitates developing novel insect management methodologies. Photostimulated behavioral responses have been implemented in a variety of model systems, such as mouse(Kim et al., 2017), zebrafish(Szobota et al., 2007; Douglass et al., 2008; Gómez-Santacana et al., 2017), *Drosophila*(Lima and Miesenböck, 2005; Alex et al., 2015), *C. elegans*(Al-Atar et al., 2009) and tadpole(Pittolo et al., 2014; Stein et al., 2012), using optogenetics or PCLs. The correlation of photoactivation of nAChRs with behavioral output is established based on DitIMI, and was successfully achieved in both transparent and opaque organisms. A majority of mosquito larvae or cockroaches harboring DitIMI exhibited fast and robust agonistic behavioral responses, indirectly demonstrating that DitIMI acts on invertebrate nAChR as an agonist. Behavioral responses occurred quickly, about 15 s after photostimulation. The results demonstrated that injected DitIMI is stable and can diffuse freely and accumulate to a sufficient level in the central neural system. Light-dependent cycles of behavioral modulation of mosquito larvae demonstrated that photoisomerization of DitIMI occurs efficiently in living organisms and the bounded closed-DitIMI in the ligand-receptor complex can be reversibly isomerized by light. Results in cockroaches illustrated that DitIMI was precisely and spatially photoactivated, however, the excitement initiated in this process cannot be recovered by visible light irradiation. This irreversibility caused the quick diffusion of active closed-DitIMI outside the light beam.

The coexistence of fluorescence and high activity of DitIMI makes it a good tracer for FP-based high-throughput screening for ligands acting on nAChRs(Allen et al., 2000). Our methodology is highly selective towards ligands sharing the same binding sites with IMI. Such nAChR fluorescence indicators have not successfully been developed probably due to the loss of activity upon large-size fluorophore attachment. Currently, radioligand binding assay is the main method for identifying ligand-protein interactions, but this technology suffers from high cost, safety issues and limited availability. Our technology provides an alternative method for nonradioactive, homogeneous and cost-effective ligand screening. Further, there is no need for separation of bound and free ligands guaranteeing undisturbed binding equilibrium, and allowing accurate quantification of binding.

## CONCLUSION

The successful development of Dit-based photochromic ligand suggested that more efforts are needed to develop Dit-derived ligands. The generation of DitIMI provides choice in the photopharmacological toolkit for elucidating functions of nAChR and may have applications for studying ligand-receptor and ligand-insect interactions. Furthermore, the establishment of FP-based screening methodology could be applied in rapid searches for potent nAChR ligands.

## MATERIALS AND METHODS

### Synthetic Procedures

The synthetic procedures and characterization of the compounds are provided in supporting information.

### Determination of photoswitching properties

The DitIMI solution (44 μM in acetonitrile) was irradiated using a UV LED (320 nm, 10 mW cm^−2^, MQK-WFH-204B, MQK, Shanghai) or blue light LED (430 nm, 5 mW cm^−2^) alternatively. Absorption spectra were recorded on a Lambda 650 UV-Vis spectrophotometer. The fatigue resistance was determined after twelve cycles of alternative irradiation of ultraviolet or blue light. The ratio of closed/open isomer (44 μM in acetonitrile) was determined by HPLC analysis before and after 320 nm light irradiation. The closed/open ratios were calculated at the isosbestic points.

### ^1^H NMR tracking

^1^H NMR spectra of DitIMI (10 mg in 0.6 mL DMSO-*d_6_*) were recorded on a Bruker AM-400 spectrometer (at 400 MHz). Then the solution was irradiated with 320 nm light at different time intervals and ^1^H NMR spectra were recorded until no changes occurred.

### Calculation of N^3^-X-N^3’^ length

The length of N^3^-X-N^3’^ of DitIMI was calculated using PyMOL 1.7.

### *In vivo* activity test

All experiments were conducted three times with three replicates in each case. Insects were considered dead if they did not move when prodded with a syringe. Two groups of DitIMI solutions in distilled water were prepared and one group was irradiated by 320 nm UV light for 15 min. Test for *Aphis craccivora*: horsebean leaves with about 50 apterous adults of *A. craccivora* were dipped in corresponding solutions for 5 s and the excess solutions were sucked out with filter paper. The insects were covered with black cloth and positioned in a temperature-controlled room (25 ± 1 °C). After a 48-hour treatment, the mortality rates were counted. Test for *Aedes albopictus*: Ten fourth-instar larvae of *A. albopictus* were transferred into plastic containers containing test solution. The containers were covered with black cloth and positioned in a temperature-controlled room. The mortality rates were calculated after a 24-hour treatment. The fluorescence images and confocal microscopic images of larvae were recorded at the same time. Test for *Periplaneta americana*: Ten cockroaches were individually treated with the test solutions by intrathoracic injection (1 μL for each cockroach) and transferred into glass jars. Then the cockroaches were covered with black cloths and positioned in a temperature-controlled room. The mortality rates were counted after a 24-hour treatment.

### Preparation of DUM neurons

Adult male cockroaches were disinfected using 75% alcohol and fixed on a wax plate. Under an anatomical lens, the cockroaches were dissected and cut along the longitudinal dorsal-median line, the back was removed, and the digestive tract, trachea, fat body and other impurities were removed and discarded. The nerve node was rinsed twice using physiological solution (185 mM NaCl, 3.0 mM KCl, 4 mM MgCl_2_, 10 mM D-glucose, 10 mM HEPES, pH = 7.2) at room temperature. The nerve nodes in physiological solution containing IA collagenase (1.5 mg mL−1) were enzymolyzed at 37 oC for 30 min. After enzymolysis, the enzymatic hydrolysate was removed, and samples were washed thrice with physiological solution (185 mM NaCl, 3.0 mM KCl, 4 mM MgCl_2_, 10 mM D-glucose, 10 mM HEPES, 5 mM CaCl_2_, fetal bovine serum (10%, V/V), 1% Penicillin-Streptomycin 100X Solution) to terminate enzymolysis. Finally, the cell suspension was filtered through a 200 mesh screen to a poly lysine coated petri dish. The cells were cultured at 28 °C for 24 h.

### Membrane potential analysis of DUM neurons

DitIMI solution (30 μM) with or without 320 nm UV light irradiation for 15 min was added to the cell suspension (1 mL). The cells were cultured at 28 °C for 30 min, washed three times with PBS and 1 mL of physiological solution (185 mM NaCl, 3.0 mM KCl, 4 mM MgCl_2_, 10 mM D-glucose, 10 mM HEPES, 5 mM CaCl_2_, fetal bovine serum (10%, V/V), 1% Penicillin-Streptomycin 100X Solution) was added. DiBAC_4_(3) (1 μL, 5 μM) was added to the cell suspension and the cells were cultured at 28 °C for 30 min and washed thrice with PBS. Finally, confocal microscopic images of DUM neurons were recorded immediately.

### Photomodulation of cockroach behaviors

Three cockroaches were fixed with pins. The first cockroach was treated with DMSO (0.4 μL) by intrathoracic injection and an optical fiber was inserted around the brain (prothoracic or mesothoracic ganglion). The second one was injected with DitIMI solution (0.4 μL, 12.5 μg/pest). The third group were injected with DitIMI solution (0.4 μL, 12.5 μg/pest) and equipped with an optical fiber around the brain (prothoracic or mesothoracic ganglion). All three groups were kept at room temperature. UV lamps (320 nm) were used as the light source. Videos of cockroaches were recorded using a camera. Each treatment had six repetitions.

### Photomodulation of mosquito larvae behaviors

The fourth-instar mosquito larvae were transferred into plastic containers containing test solution (4 mL) and were cultured at 25 °C for 6 h. Three sets of experiments were set up (two fourth-instar mosquito larvae for each group). The first group of larvae treated with 5% DMSO aqueous solution was irradiated with 320 nm UV light. The second group of larvae treated with DitIMI (30 μM) were cultured in the dark. The third group of larvae treated with DitIMI (30 μM) were alternatively irradiated by 320 nm UV light or green light (520 nm) for 90s. Videos of larvae were recorded by a super digital microscope (Chengdu Liyang, China). Each treatment had six repetitions.

### Fluorescence polarization analysis

The ventral nerve-cord membranes were prepared according to a modified reported procedure (Miyagi et al., 2006). American cockroaches were dissected in ice-cold 50 mM Tris-HCl buffer containing 200 mM sucrose and 1 mM EDTA (pH=7.4, buffer A). The isolated nerve cords were homogenized on ice with a glass-Teflon homogenizer (25 strokes) in 50 mM Tris-HCl (pH = 7.4), 200 mM sucrose and 1 mM EDTA. The homogenate was centrifuged at 25,000 g for 30 min. The supernatant was decanted, and the pellet was washed with buffer (2 mL) twice. The pellet was resuspended in ice-cold 50 mM Tris-HCl, 120 mM NaCl and stored at −80 °C. The protein-dye binding method was used to determine the protein concentration with bovine serum albumin (BSA) as a standard. The wavelengths for excitation and emission were 420 and 590 nm, respectively. The mixture of nAChR membranes (99 μL, 227 mg/L) and varying concentrations of DitIMI (1 μL, 1.95, 3.91, 7.81, 15.625, 31.25, 62.50 and 80 μM, irradiated at 320 nm UV light for 15 min) in black, low-binding half area 96-well plates (Corning 3993) was incubated at 22 °C for 1 h. The FP signals were recorded three times using a Synergy H1 microplate reader (Bio-Tek, Winooski, VT, USA) to give the binding curve. For the competitive binding assay, a mixture of nAChR membranes (98 μL, 232 mg/L), DitIMI (1 μL, 4 mM, irradiated at 320 nm UV light for 15 min) and nAChR ligands (1 μL, 0.001, 0.003, 0.01, 0.03, 0.1, 0.3, 1, 3 and 10 μM) was incubated at 22 °C for 1 h and the FP signals were then recorded. The saturation binding and percentage inhibition were determined from three separate experiments, each with triplicate samples.

## ACKNOWLEDGEMENTS

We thank Professor John E. Casida of University of California at Berkeley for facilitating nAChR binding assays *in vitro* and for help in manuscript preparation. This work was financially supported by the National Key Research and Development Program of China (2018YFD0200100), the National Natural Science Foundation of China (No. 21877039, 32072441) and the Innovation Program of Shanghai Municipal Education Commission (2017-01-07-00-02-E00037).

---

**Figure 1.**
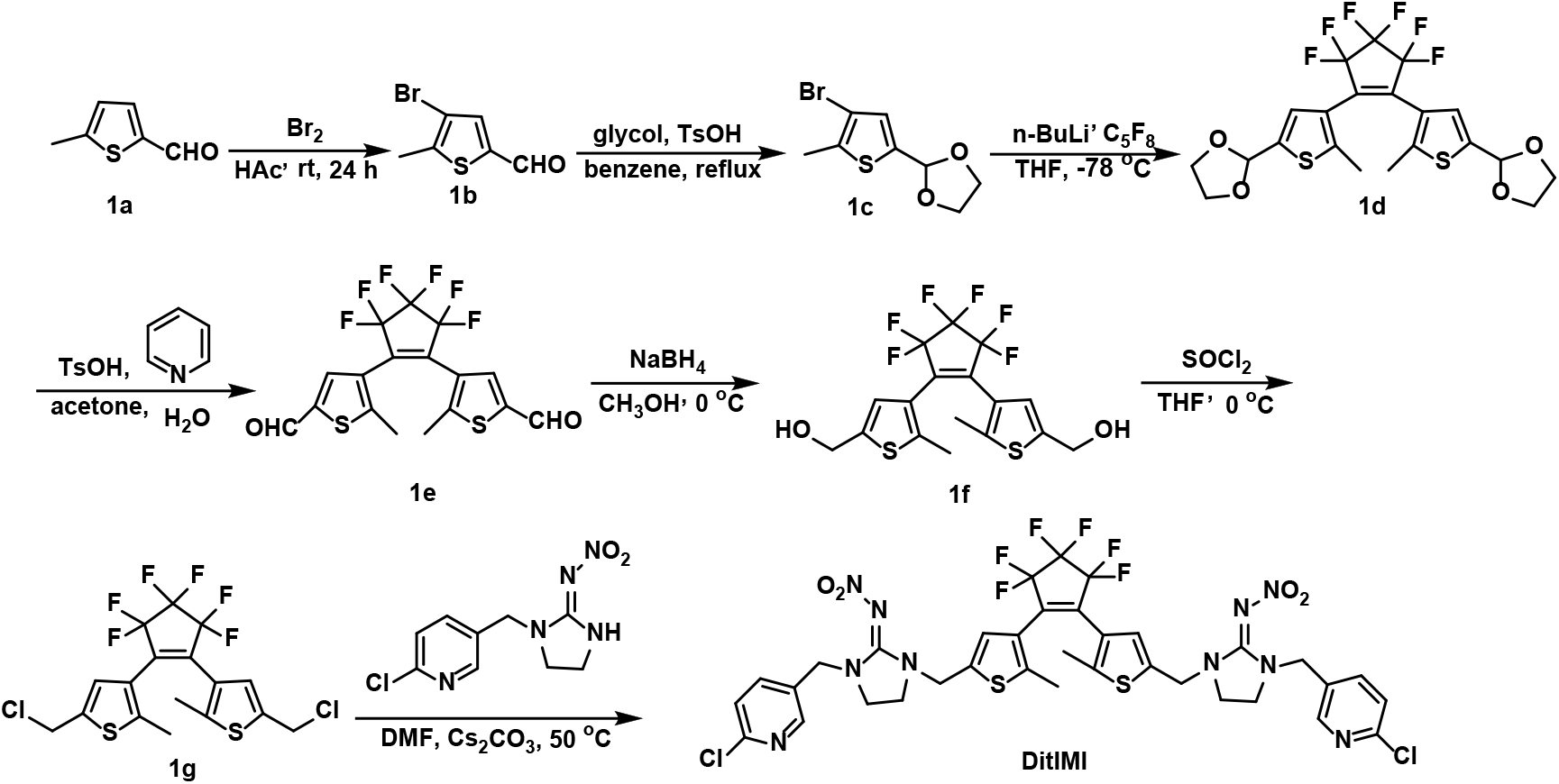
Synthetic routes of DitIMI.

**Figure 2.**
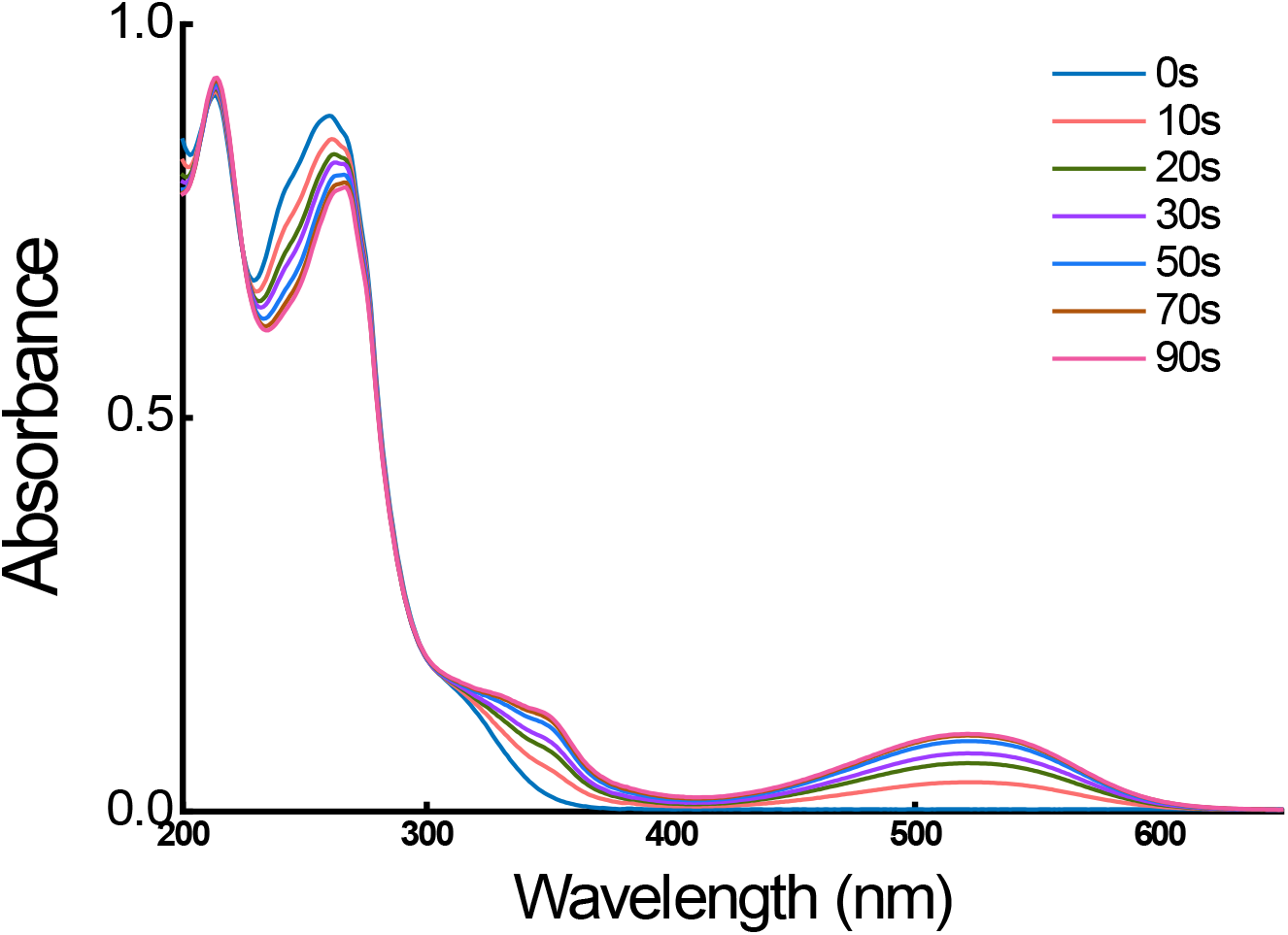
UV-Vis absorbance of DitIMI under UV irradiation at 320 nm at varying times (1-90 s).

**Figure 3.**
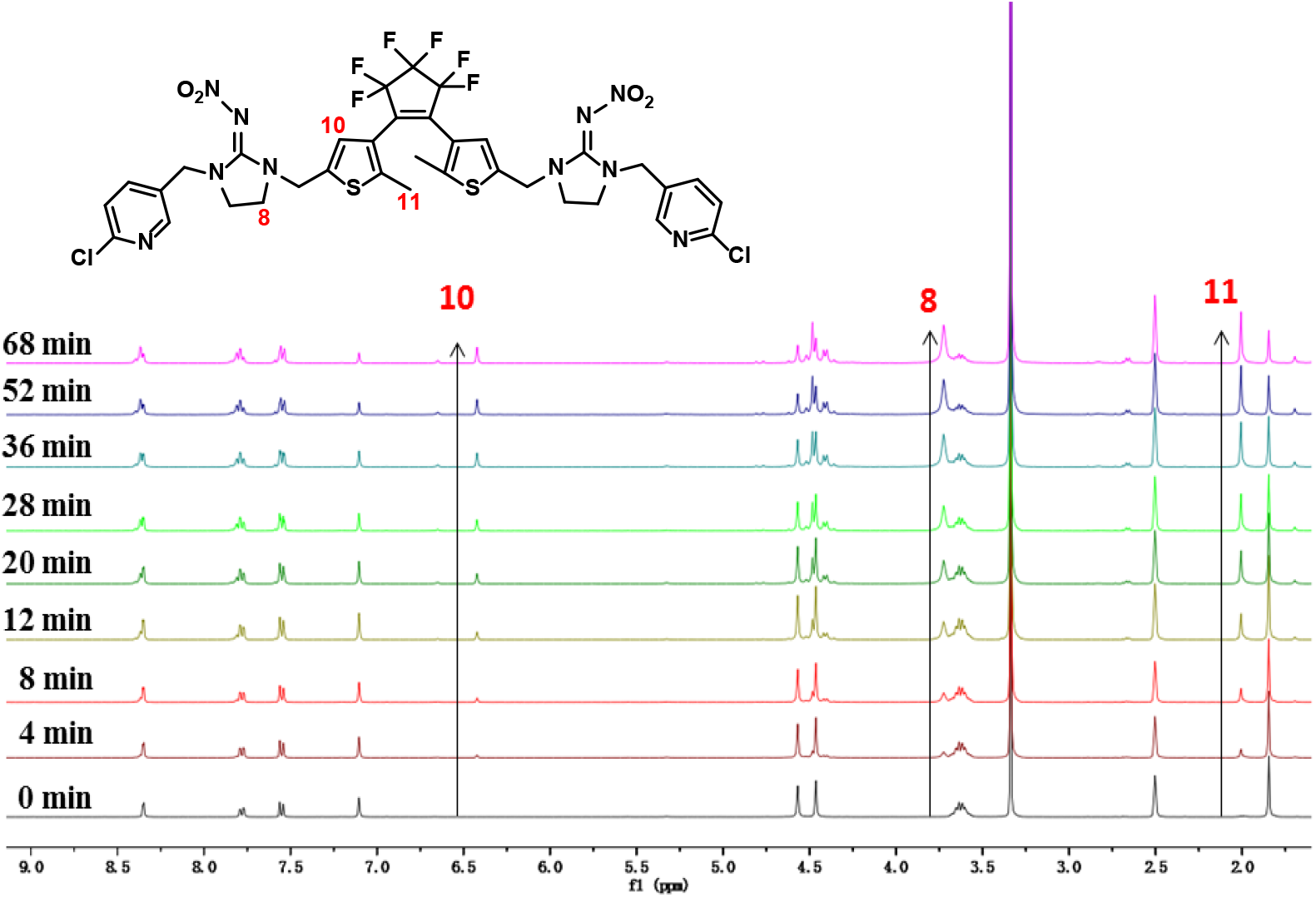
^1^H NMR spectral changes of DitIMI in DMSO-*d_6_* upon irradiation with UV light at 320 nm.

**Figure 4.**
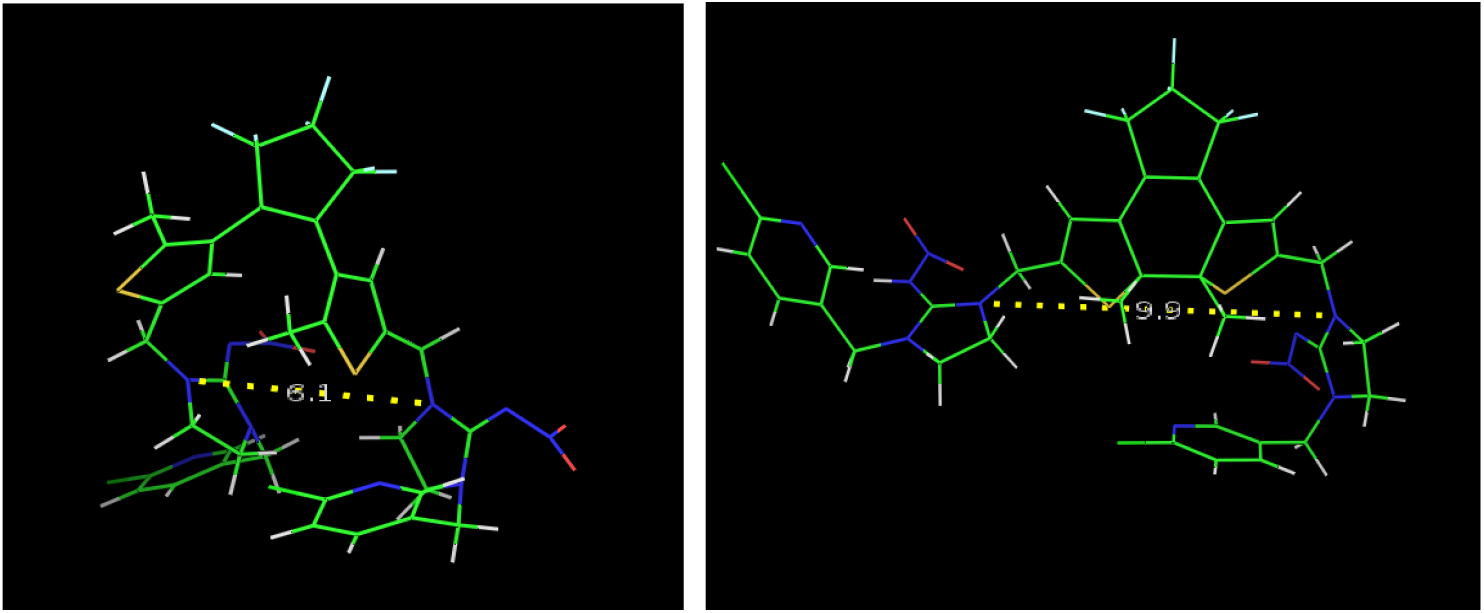
The length of N^3^-X-N^3’^ of both photoisomers of DitIMI.

**Figure 5.**
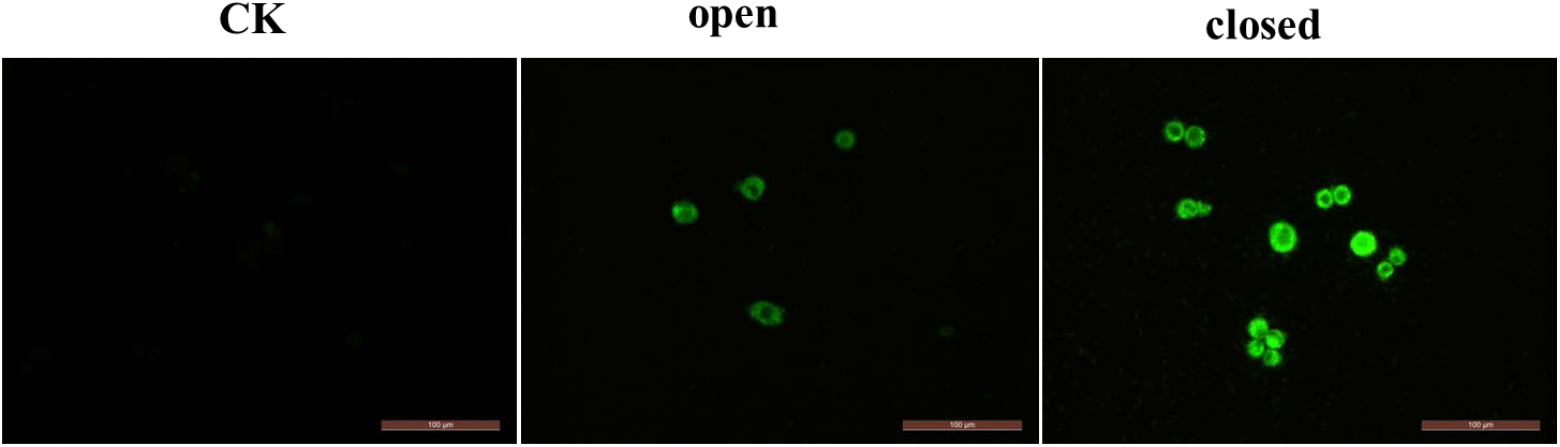
The membrane potential influence of insect Sf9 cells of DitIMI using DiBAC4(3) as a fluorescence indicator.

**Figure 6.**
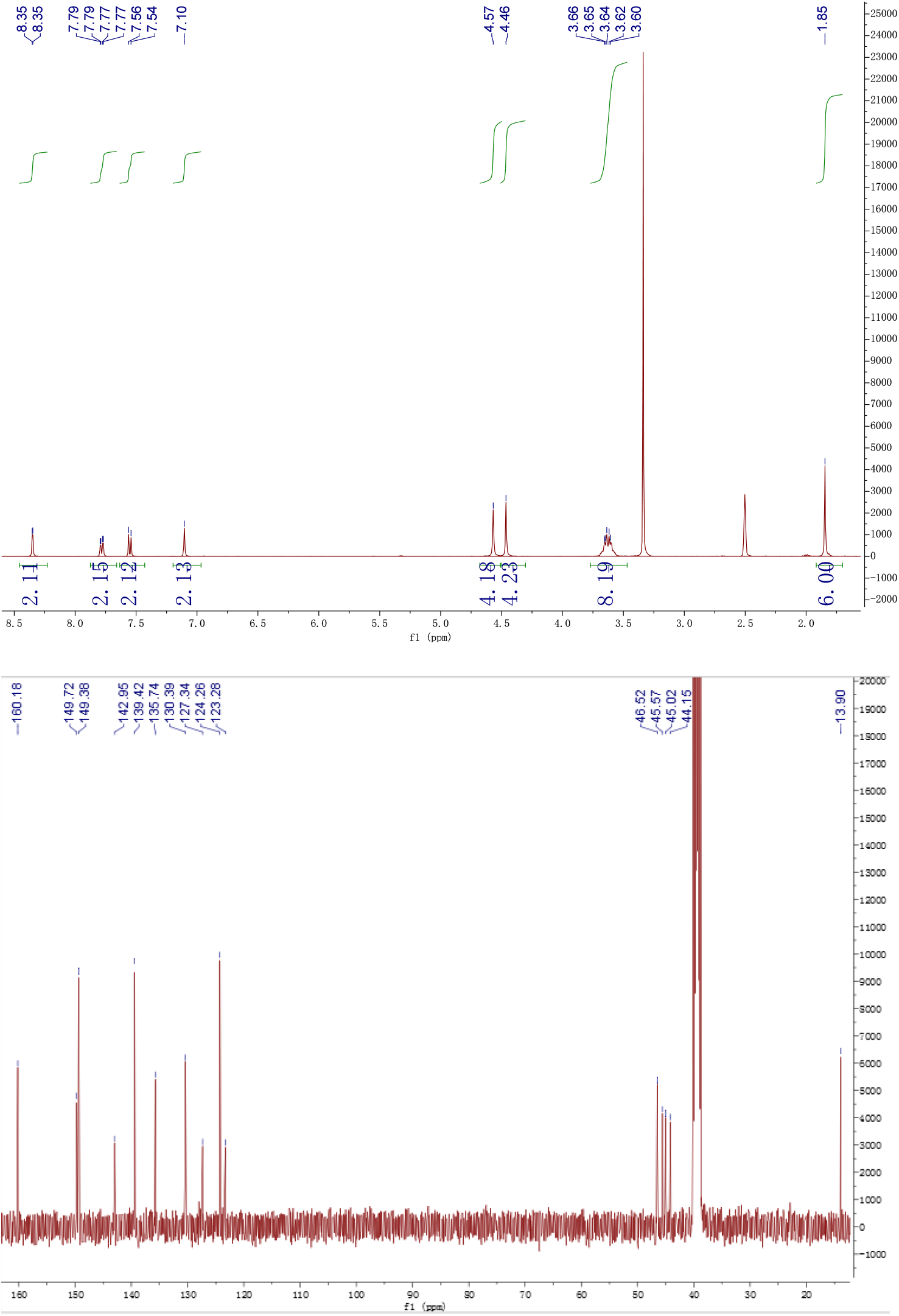
^1^H NMR and ^13^C NMR spectra of DitIMI.

**Figure 7.**
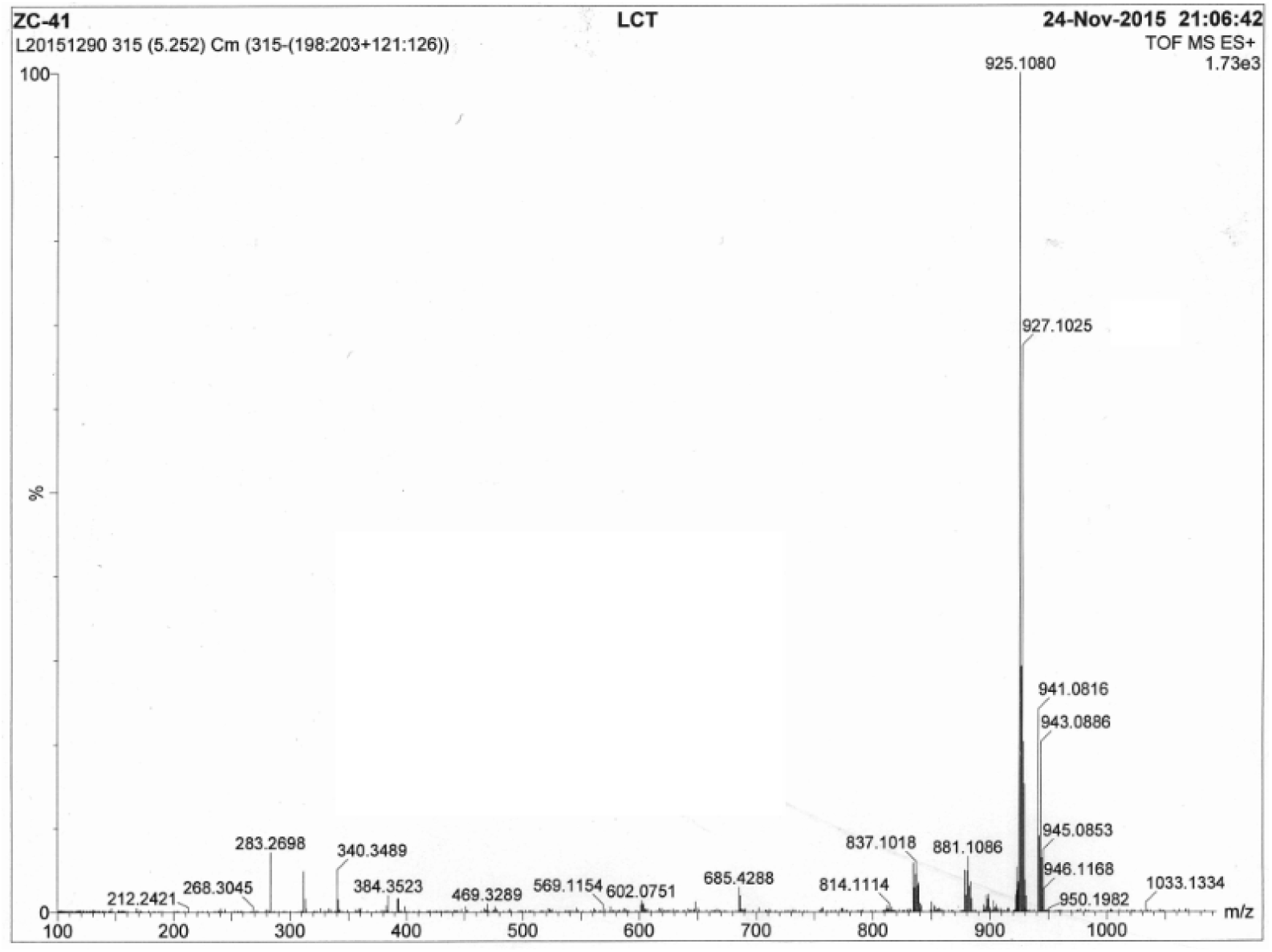
HRMS spectrum of DitIMI.

**Figure 8.**
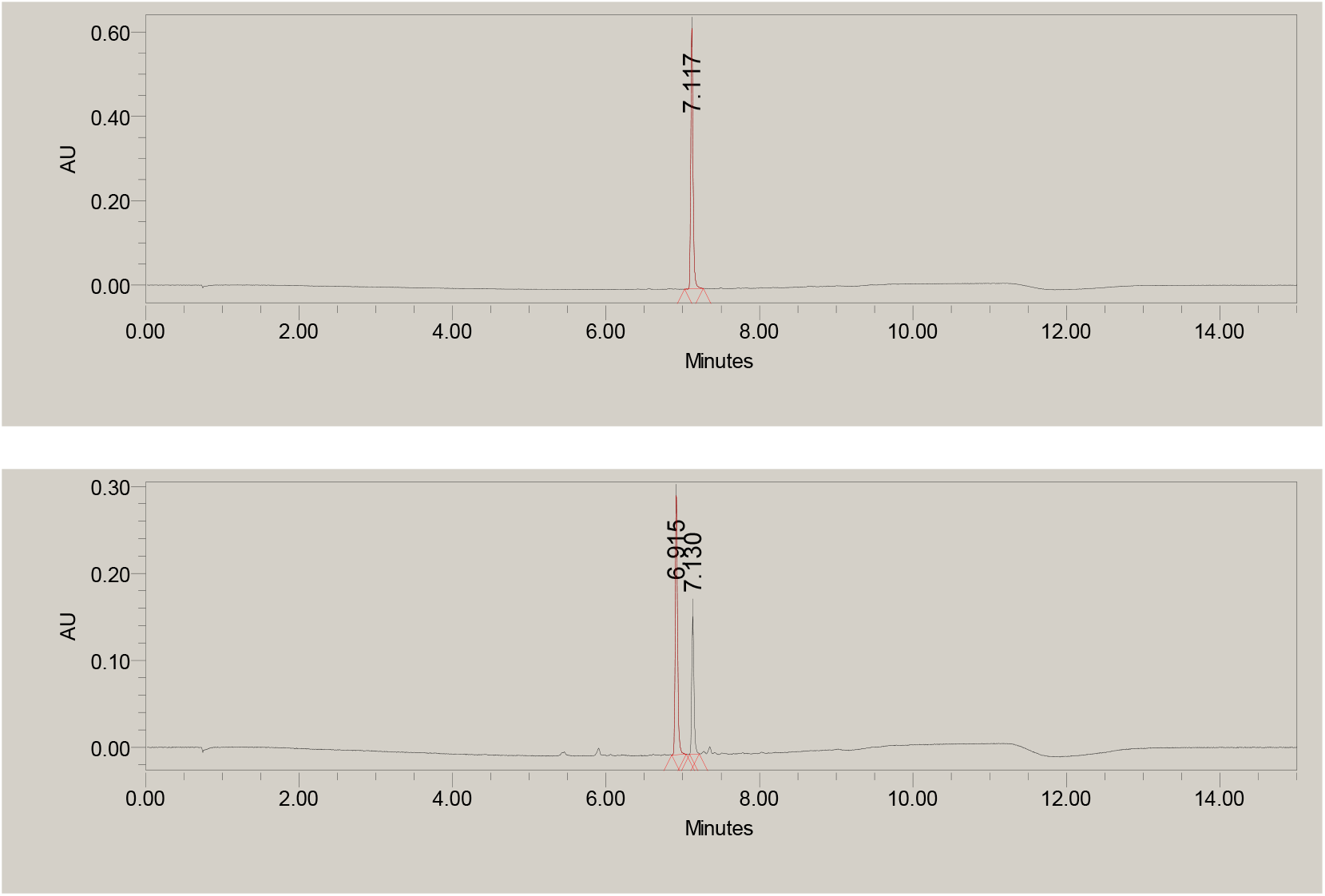
HPLC spectrum of DitIMI before and after UV irradiation.

**Table 1.** Maximum absorbance wavelength (λ max, nm), ratio of photoisomers of DitIMIs.

| Compd. | $\lambda_{\max} \pi-\pi^*$ | nonirradiated | $\lambda_{\max} \pi-\pi^*$ | irradiated |
| --- | --- | --- | --- | --- |
|  | (open, nm) | open: close | (close, nm) | open: close |
| DitlIMI | 310 | 100:0 | 520 | 33:67 |

**Table 2.** Physicochemical properties of DitIMI and IMI.

| Compd. | LogP |  | LogS |  | PKa |  | TPSA (Å<sup>2</sup>) d |  |
| --- | --- | --- | --- | --- | --- | --- | --- | --- |
|  | open | closed | open | closed | open | closed | open | closed |
| DitlIMI | 7.29 | 8.34 | -12.33 | -10.65 | 8.06 | 9.41 | 166.02 | 166.02 |
| IMI | 1.81 |  | -2.43 |  | 11.71 |  | 91.80 |  |

**Table 3.** Behavioral response of mosquito larvae (*Aedes albopictus*) treated with irradiated or unirradiated DitIMI.

| DitlIMI (μM) | Behavioral responses of *Aphis craccivora* after 2 h treatment |  |
| --- | --- | --- |
|  | nonirradiated | irradiated |
| 440 | + | ++ |
| 220 | + | ++ |
| 110 | + | ++ |
| 55 | - | ++ |
| 25 | - | ++ |
| 12.5 | - | ++ |
| 6.25 | - | + |
| 3.13 | - | + |
| 1.63 | - | + |
| CK | - | - |

Note: -, no excited response was observed; +, part of *Aedes* individual showed excitement; ++, entire *Aedes* individual showed excitement.

**Table 4.** Behavioral responses of American cockroach (*P. americana*) treated with irradiated or unirradiated DitIMI.

| DitIMI (µg/pest) | Behavioral responses of *P. americana* after 1 h treatment |  |
| --- | --- | --- |
|  | nonirradiated | irradiated |
| 400 | ++ | ++ |
| 200 | ++ | ++ |
| 100 | + | ++ |
| 50 | + | ++ |
| 25 | + | ++ |
| 12.5 | - | + |
| 6.25 | - | - |
| CK | - | - |

Note: -, no excited response was observed; +, part of *Periplaneta* showed excitement; ++, entire *Periplaneta* showed excitement.

## Instruments and Chemicals

Melting points were determined on a Büchi melting point B-540 apparatus (Büchi Labortechnik AG, Flawil, Switzerland). ^1^H NMR and ^13^C NMR spectra were recorded on a Bruker AM-400 spectrometer (at 400 MHz and 100 MHz for ^1^H NMR and ^13^C NMR, respectively) (Bruker, Karlsruhe, Germany) with CDCl_3_ or DMSO-*d_6_* as the solvent and TMS as the internal standard. Chemical shifts are reported in *δ* (parts per million) values and coupling constants (*J*) are denoted in Hz. High-resolution electron mass spectra (ESI-TOF) were obtained using a Micromass LC-TOF spectrometer. Column chromatography was performed using silica gel (200-300 mesh, Hailang, Qingdao). The UV-Vis spectra and fatigue resistance were obtained on a Lambda 650 UV-Vis spectrophotometer (PerkinElmer, Inc., Fremont, California, USA). Chromatographic analysis was performed using an ACQUITY UPLC-H Class system (Waters Corp., USA) equipped with HSS T3 reversed phase column with 100 mm × 2.1 mm i.d. and 1.8 μm particle size, a quaternary solvent delivery system, a 48-vial autosampler (10 μL loop) and a photodiode array detector (PDA). Fluorescence emission spectra were recorded on a fluorescence spectrophotometer (Varian, Palo Alto, California, USA). Fluorescence images were taken on a Leica DMI3000B Fluorescent Microscope (Leica Microsystems Wetzlar GmbH, Wetzlar, Germany). Confocal microscopic images were recorded using a Nikon-A1R Confocal Laser Microscope (Nikon Co., Tokyo, Japan), and microscope images were recorded on a LY-WN-YC600 digital microscope (Liyang Precision Machinery Co. Ltd., Chengdu, China). Fluorescence polarization was carried out on a Synergy H1 microplate reader (Bio-Tek, Winooski, VT, USA). Reagents and solvents were used as received from commercial suppliers. Yields were not optimized.

### Insects

The 4th-instar mosquito larvae (*Aedes albopictus*) were obtained from the National South Pesticide Initiative Center in Shanghai, China. The armyworm (*Mythimna separata*) and aphid (*Aphis craccivora*) were obtained from Shanghai Key Laboratory of Chemical Biology, East China University of Science and Technology in Shanghai, China. Cockroaches (*Periplaneta americana*) were obtained from Flying Pharmaceutical Co., Ltd. in Danyang, China.

### Synthetic procedures of DitIMI

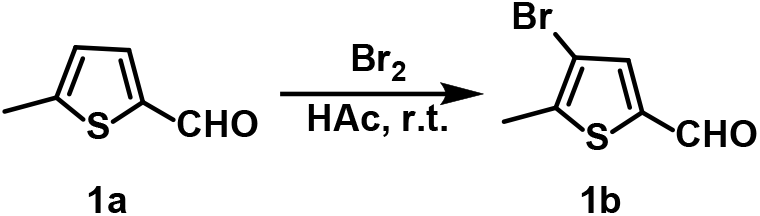
4-bromo-5-methyl-2-formylthiophene (**1b**)

To a stirred solution of 5-methyl-2-formylthiophene (**1a**) (6.3 g, 50.0 mmol) in acetic acid was added an acetic solution of Br_2_ (8.78 g, 55.0 mmol) at room temperature. Stirring was continued for 24 h and the reaction was stopped by the addition of water. The reaction mixture was neutralized by Na_2_CO_3_ to neutrality. The mixture was extracted with ether, dried and filtrated. The solvent was evaporated to obtain the black solid. The residue was purified by chromatography on a silica gel with petroleum ether/ethyl acetate (v/v = 40/3) as eluent to afford **1b** (3.1 g) in 30.1% yield. ^1^H NMR (400 MHz, CDCl_3_) *δ* 9.78 (s, 1H), 7.26 (s, 1H), 2.49 (s, 3H). ^13^C NMR (101 MHz, CDCl_3_) *δ* 181.60, 145.84, 140.12, 138.73, 111.24, 15.93.

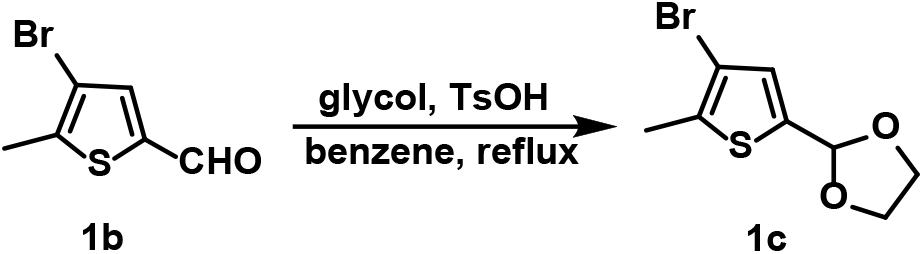
4-bromo-5-methyl-2-(1,3-dioxolane)thiophene (**1c**)

4-Bromo-5-methyl-2-formylthiophene (**1b**) (2.0 g, 9.75 mmol), glycol (1.23 g, 19.51 mmol) and p-toluenesulfonic acid (0.15 g, 97.53 mmol) were dissolved in benzene. The reaction was refluxed for 4 h and then washed sequentially three times with NaOH (3.0 mol/L) and water (15 mL × 3). The combined benzene layers were dried over Na_2_SO_4_ and the filtrate was concentrated under reduced pressure. The residue was then chromatographed on a silica gel with petroleum ether/ethyl acetate (v/v = 20/1) as eluent to afford **1c** (2.4 g) in 90.8% yield. ^1^H NMR (400 MHz, CDCl_3_) *δ* 6.96 (s, 1H), 6.00 (s, 1H), 4.18-3.88 (m, 4H), 2.38 (s, 3H). ^13^C NMR (101 MHz, CDCl_3_) *δ* 138.70, 135.35, 128.79, 108.45, 99.69, 65.19, 14.86.

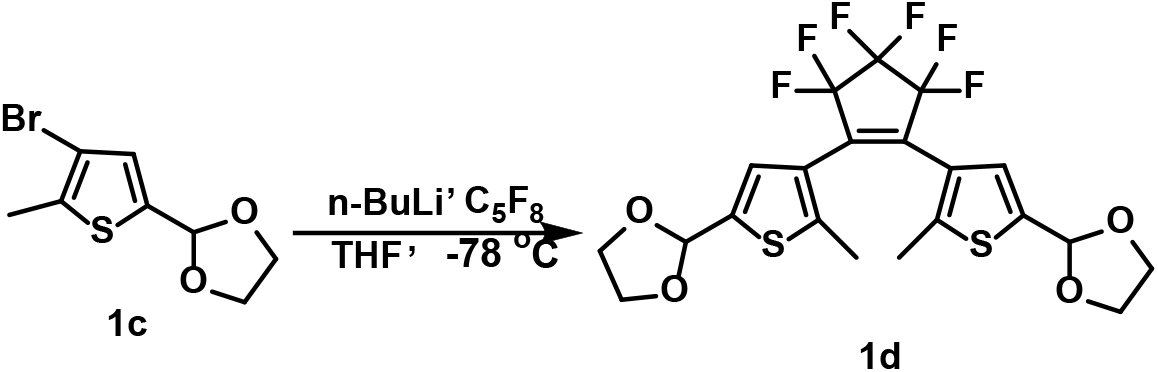
1,2-Bis[2-methyl-5-(1,3-dioxolane)-3-thienyl]perfluorocyclopentene (**1d**)

To a stirred solution of **1c** (2.4 g, 9.63 mmol) in dry THF was added 2.5 mol/L n-BuLi/hexane (7.7 mL, 19.27 mmol) at −78 °C under a nitrogen atmosphere. After stirring for 40 min, C_5_F_8_ (1.23 g, 5.78 mmol) was slowly added to the reaction mixture and stirred for 2 h at this temperature. The reaction was quenched with water and extracted with ether. The organic layer was collected and washed with water (15 mL × 3). The organic layer was dried, filtered and evaporated resulting in **1d** (2.0 g) which was used directly with our purification.

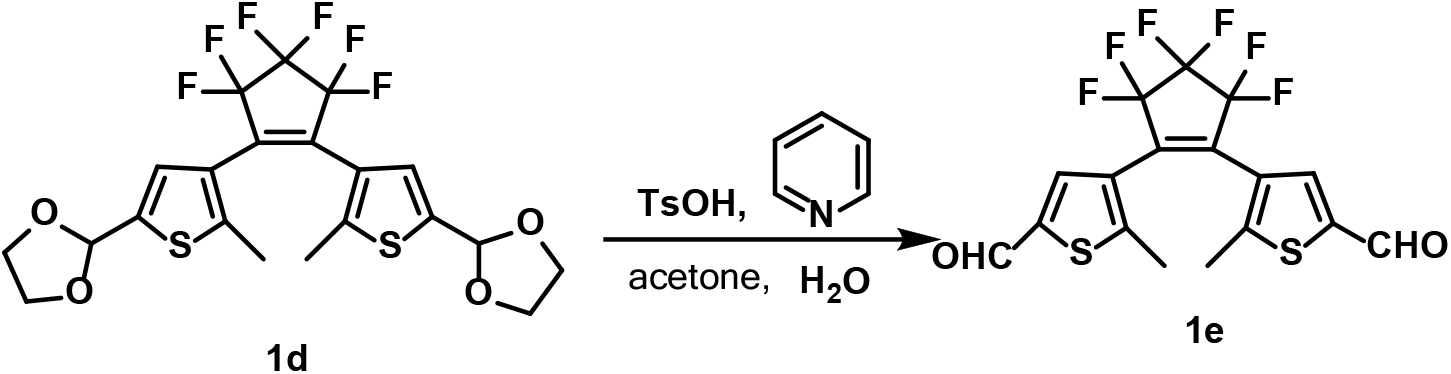
1,2-Bis(2-methyl-5-formyl-3-thienyl)perfluorocyclopentene (**1e**)

The raw **1d** (2.0 g, 3.90 mmol), p-toluenesulfonic acid (0.4 g, 2.22 mmol) and pyridine (2.0 mL) in a mixture of water (30 mL) and acetone (80 mL) was refluxed for 24 h. After completion, the reaction mixture was washed sequentially by aqueous NaHCO3 and water (25 mL × 3). The organic layer was dried over anhydrous Na_2_SO_4_, filtrated and evaporated. The crude product was purified by column chromatography on SiO_2_ using ethyl acetate and petroleum ether mixture (v/v = 1/5) as the eluent affording **1e** (0.15 g) in 87% yield. ^1^H NMR (400 MHz, CDCl_3_) *δ* 9.86 (s, 2H), 7.74 (s, 2H), 2.03 (s, 6H). ^19^F NMR (376 MHz, CDCl_3_) δ −110.30 (t, *J* = 5.1 Hz, 4F), −130.57 — −133.99 (m, 2F).

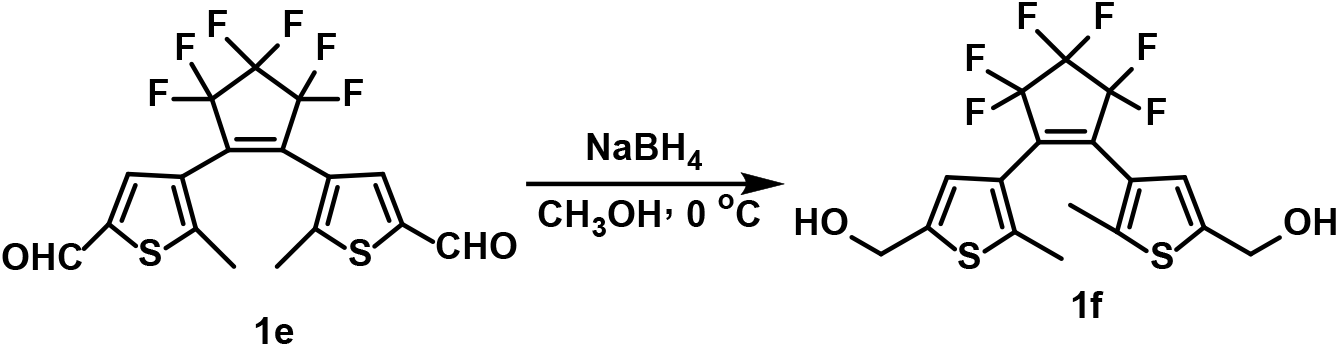
1,2-Bis(2-methyl-5-methylol-3-thienyl)perfluorocyclopentene (**1f**)

To a solution of **1e** (0.45 g, 1.06 mmol) in methanol (35 mL) was slowly added a sodium borohydride (0.16 g, 4.24 mmol) in methanol (7 mL) and water (7 mL). After stirring for 1 h at 0 °C temperature, the reaction mixture was extracted with ether (30 mL × 4), and the combined organic layer was washed with saturated aqueous brine (15 mL × 3) and dried over MgSO_4_. Removal of the solvent gave **1f** as a yellow solid. The crude product was purified by column chromatography on SiO_2_ using ethyl acetate and petroleum ether mixture (v/v=1/1) as the eluent affording **1f** (0.45 g) in 99% yield. 1H NMR (400 MHz, CDCl3) *δ* 6.95 (s, 2H), 4.76 (s, 4H), 1.88 (s, 8H). ^19^F NMR (376 MHz, CDCl_3_) δ −110.10 (t, *J* = 10.9 Hz, 4F), −131.48 — −134.27 (m, 2F).

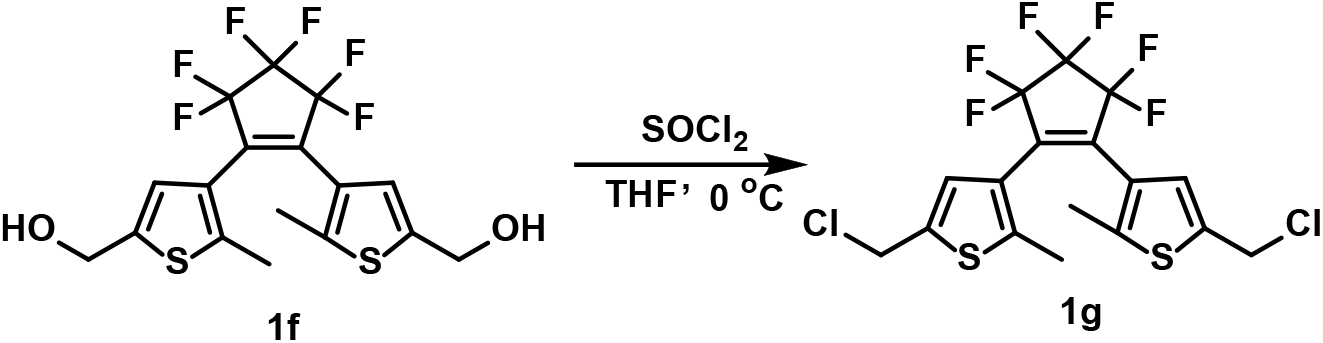
1,2-Bis(2-methyl-5-chloromethyl-3-thienyl)perfluorocyclopentene (**1g**)

To a mixture of **1f** (0.25 g, 0.58 mmol), dry pyridine (1.0 mL) and dry THF (40 mL) was added thionyl chloride (0.28 g, 2.33 mmol) at 0 °C under argon. The reaction mixture was stirred for 2 h at 0 °C. After completion, the mixture was poured into ice water (40 mL) and extracted with ether (20 mL × 4). The combined organic phases were washed with saturated NaHCO_3_ (30 mL), dried over anhydrous MgSO_4_ and evaporated *in vacuo* to give **1g** as a brown solid (0.25 g) in 92% yield, which was used for the next step immediately without further purification.

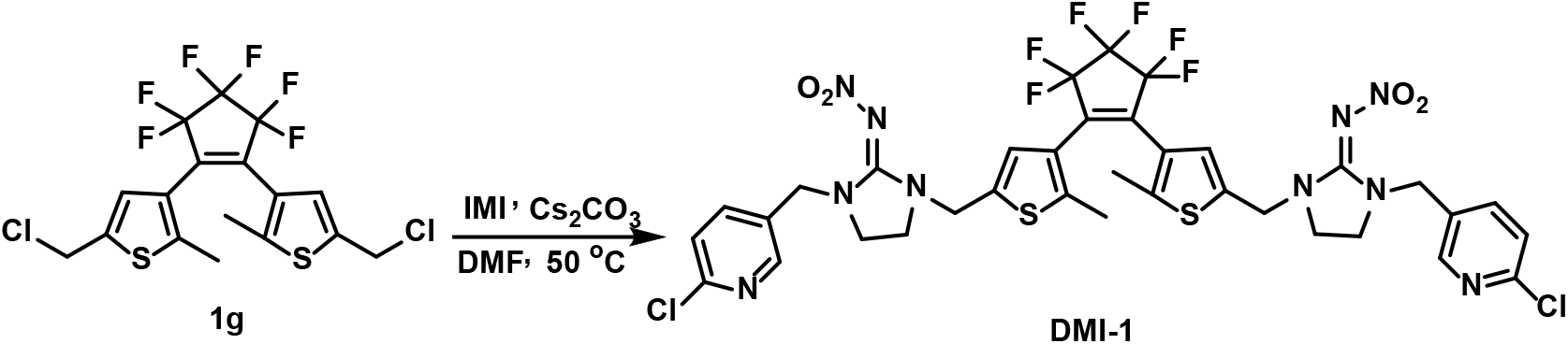

### DitIMI

Imidacloprid (IMI) (0.40 g, 1.55 mmol), Cs_2_CO_3_ (0.72 g, 2.19 mmol) and DMF (5 mL) were stirred under nitrogen at 50 °C for 0.5 h, then **1g** (0.30 g, 0.65 mmol) was added to the mixture. The resulting mixture was stirred under nitrogen at 50 °C for 4 h. The precipitate was separated by filtration. The solvent was removed, and the crude product was purified by column chromatography on SiO_2_ using dichloromethane and methanol mixture (v/v=20/1) as the eluent affording the target compound DitIMI (0.35 g) in 60% yield. m.p. 108.7-110.9 °C. ^1^H NMR (400 MHz, DMSO-*d6*) *δ* 8.35 (d, *J* = 2.0 Hz, 2H), 7.78 (dd, *J* = 8.3, 2.4 Hz, 2H), 7.55 (d, *J* = 8.2 Hz, 2H), 7.10 (s, 2H), 4.53 (d, *J* = 28.1 Hz, 4H), 4.46 (s, 4H), 3.77− 3.47 (m, 8H), 1.83 (d, *J* = 13.4 Hz, 6H). ^13^C NMR (101 MHz, DMSO-*d_6_*) δ 160.19, 149.72, 149.53, 149.38, 142.96, 139.42, 135.74, 130.39, 127.34, 123.28, 46.52, 45.57, 45.02, 44.15, 13.90. ^19^F NMR (376 MHz, DMSO-*d_6_*) *δ* −109.35 (t, *J* = 10.9 Hz, 4F), −131.11 (m, 4F). HRMS (ESI-TOF): m/z calcd for C_35_H_30_^35^Cl_2_F_6_N_10_O_4_S_2_Na [M+Na]^+^ 925.1072, found 925.1080; m/z calcd for C_35_H_30_^35^Cl^37^ClF_6_N_10_O_4_S_2_Na [M+Na]^+^ 927.1042, found 927.1025; m/z calcd for C_35_H_30_^37^Cl_2_F_6_N_10_O_4_S_2_Na [M + Na]^+^ 929.1013, found 929.1030.

